# Neurofunctional basis underlying audiovisual integration of print and speech sound in Chinese children

**DOI:** 10.1101/2020.05.31.126128

**Authors:** Zhichao Xia, Ting Yang, Xin Cui, Fumiko Hoeft, Hong Liu, Xianglin Zhang, Hua Shu, Xiangping Liu

**Affiliations:** State Key Laboratory of Cognitive Neuroscience and Learning & IDG/McGovern Institute for Brain Research, Beijing Normal University, China; School of Systems Science, Beijing Normal University, China; Faculty of Psychology, Beijing Normal University, China; Beijing Key Laboratory of Applied Experimental Psychology, National Demonstration Center for Experimental Psychology Education, Faculty of Psychology, Beijing Normal University, China; Haskins Laboratories, USA; Department of Psychological Sciences and Brain Imaging Research Center, University of Connecticut, USA; Department of Psychiatry and Weill Institute for Neurosciences and Dyslexia Center, University of California, San Francisco, USA; Department of Neuropsychiatry, Keio University School of Medicine, Japan

**Keywords:** audiovisual integration, character, fMRI, pinyin, Chinese, reading development

## Abstract

Effortless print-sound integration is essential to reading development, and the superior temporal cortex (STC) is the most critical brain region. However, to date, the conclusion is almost restricted to alphabetic orthographies. To examine the neural basis in non-alphabetic languages and its relationship with reading abilities, we conducted a functional magnetic resonance imaging study in typically developing Chinese children. Two neuroimaging-based indicators of audiovisual processing—additive enhancement (higher activation in the congruent than the average activation of unimodal conditions) and neural integration (different activations between the congruent versus incongruent conditions)—were used to investigate character-sounds (opaque) and pinyin-sounds (transparent) processing. We found additive enhancement in bilateral STCs processing both character and pinyin stimulations. Moreover, the neural integrations in the left STC for the two scripts were strongly correlated. In terms of differentiation, first, areas beyond the STCs showed additive enhancement in processing pinyin-sounds. Second, while the bilateral STCs, left inferior/middle frontal and parietal regions manifested a striking neural integration (incongruent > congruent) for character-sounds, no significant clusters were revealed for pinyin-sounds. Finally, the neural integration in the left middle frontal gyrus for characters was specifically associated with silent reading comprehension proficiency, indicating automatic semantic processing during implicit character-sound integration. In contrast, the neural integration in the left STC for pinyin was specifically associated with oral reading fluency that relies on grapho-phonological mapping. To summarize, this study revealed both script-universal and script-specific neurofunctional substrates of print-sound integration as well as their processing- and region-dependent associations with reading abilities in typical Chinese children.

**Highlights:**

- The bilateral superior temporal cortices display additive enhancement in processing print-sound information regardless of orthographies.
- The location, direction, and strength of the neural audiovisual integration are modulated by orthographic transparency.
- The neural integration in the left middle frontal gyrus in processing character-sounds is correlated to reading comprehension.
- The neural integration in the left superior temporal gyrus for pinyin is correlated with oral reading fluency.
- The neural integrations of character-sounds and pinyin-sounds show spatial overlap and inter-subject correlation in the left superior temporal cortex.

## Introduction

Efficient print-speech sound mapping is a prerequisite for successful reading acquisition and a hallmark of fluent reading (Blomert & Froyen, 2010; Richlan, 2019). Better knowledge about the underlying neural basis can further our understanding of how this process contributes to reading development and provide objective training targets for children with reading disorders (RD). With neuroimaging-based indicators such as audiovisual additive enhancement and neural integration, studies revealed that the automatic print-sound integration uniquely accounts for reading outcomes in both shallow (e.g., Dutch, German) and opaque orthographies (e.g., English) (Bakos, Landerl, Bartling, Schulte-Körne, & Moll, 2017; Blau et al., 2010; Blomert & Willems, 2010; Kronschnabel, Brem, Maurer, & Brandeis, 2014; McNorgan, Wagner, & Booth, 2013), while the bilateral superior temporal cortices (STCs) as the most important brain area (Blomert, 2011; van Atteveldt & Ansari, 2014). However, the current evidence is almost restricted to alphabetic languages. The question of whether the same underlying neural substrates exist and play a critical role in reading in non-alphabetic languages with dramatically different linguistic characteristics remains largely open.

Chinese, a logographic writing system, is particularly suited for answering this cross-linguistic question given its extremely deep orthography (Perfetti, Cao, & Booth, 2013). The print-sound mapping is arbitrary given that Chinese characters directly map to syllables and there are no grapheme-phoneme correspondence (GPC) rules. Moreover, in many cases, different characters share the same pronunciation (i.e., homophones), whereas a single character can also have multiple sounds (i.e., polyphone). Therefore, the link between print (character) and speech sound (syllable) is even more opaque. Of importance, in comparison with grapho-phonological mapping, grapho-semantic correspondence is more systematic in Chinese characters, deeply involved in character recognition, and plays a role equal to or even more essential than grapho-phonological processing in reading acquisition (Liu et al., 2017; Ruan, Georgiou, Song, Li, & Shu, 2018; Yang, Shu, McCandliss, & Zevin, 2013; Yang, Zevin, Shu, McCandliss, & Li, 2006). These linguistic characteristics could bring language-specific neural features accompanying reading-related processes.

Recently, Xu et al. (2019) conducted a magnetoencephalography (MEG) study in Chinese adults and identified that the left frontal cortex was also involved in processing character-sound information besides the left STC. However, some limitations need to be noted. First, since no behavior measures were included, the functional role of the significant effect can only be speculated without evidence. Moreover, for experienced adult readers, audiovisual integration of characters and corresponding speech sounds is fully automated. Thus, an important but unanswered question is about the neurofunctional features of this process in the developing brain. Against this background, the current study was to investigate the neural basis of character-sound integration and its association with reading abilities in typically developing Chinese children (aim 1).

In particular, we focused on the early-acquired single-element pictographic characters given their fundamental role in learning-to-read in Chinese (Shu, Chen, Anderson, Wu, & Xuan, 2003). At the very beginning stage, a small set of characters are taught first. These simple characters will later be used as radicals in complex compound characters (i.e., phonograms), which comprise greater than 80% of Chinese characters (Chan & Siegel, 2001). Importantly, as a pictographic character does not contain any components (i.e., radicals), there is no clue on its pronunciation or meaning. Thus, the mappings between its visual form with sound (syllable) and meaning (morpheme) are entirely arbitrary.

It should be noted that children in Mainland China and Singapore learn pronunciations of characters with Hanyu pinyin (pinyin hereafter). In brief, pinyin is an official Roman alphabetic system used to teach the sounds of characters (Yin et al., 2011). A child learns pinyin immediately after entering school and then uses it to assist in conquering the pronunciation of new characters (Guan, Liu, Chan, Ye, & Perfetti, 2011; Shu, Peng, & McBride-Chang, 2008; Yan, Miller, Li, & Shu, 2008). In other words, while conquering the sounds of letters to an automatic level is one of the most critical prerequisites in alphabetic languages (Blomert, 2011; Richlan, 2019), learning pinyin plays a scaffolding role at the initial stage of Chinese reading acquisition. Pinyin borrows letters from alphabets, and most have consistent mappings with speech sounds (phonemes), so it has a transparent orthography. Therefore, we included pinyin in this study to further examine whether and how the neural features of print-sound integration are associated with orthographic depth and provide information that helps illustrate the role of pinyin in developing fluent reading ability in Chinese (aim 2).

### Present study

In this functional magnetic resonance imaging (fMRI) study, we planned to use two indicators—additive enhancement and neural integration—to identify brain regions involved in different aspects of multimodal information processing. Notably, additive enhancement and neural integration could be observed in passive paradigms, pinpointing an automatic nature (Blau et al., 2010; van Atteveldt, Formisano, Goebel, & Blomert, 2007).

The additive enhancement is based on the additive model that was initially examined in animal electrophysiological studies where researchers found audiovisual signals induced stronger (i.e., super-additive) or lower (i.e., sub-additive) neural response than the summation of responses to unimodal auditory and visual inputs in areas such as the STC (Stein & Stanford, 2008). However, the super-additive enhancement criterion is too strict about detecting multisensory integration regions with fMRI data. In contrast, with liberal variants such as mean additivity enhancement criterion (congruent condition > mean of unimodal auditory and visual conditions), areas including the bilateral STCs can be successfully identified (Beauchamp, 2005; Kronschnabel et al., 2014; McNorgan & Booth, 2015). In terms of functional significance, the super-(and mean) additive enhancement occurs early and is more likely to reflect more general and fundamental aspects of multisensory processing (Kronschnabel et al., 2014; Xu et al., 2019). In line with it, this effect has been observed in both shallow (e.g., German) (Kronschnabel et al., 2014) and opaque (e.g., English) (McNorgan & Booth, 2015) alphabetic orthographies and no associations between additive enhancement and reading performance were found (Kronschnabel et al., 2014). In this case, we expected that the bilateral STCs would show an additive enhancement both in processing character-related and pinyin-related audiovisual information.

The neural integration—different brain response to audiovisual congruent versus incongruent pairs—is another widely-used indicator of multisensory integration and repeatedly reported in the left STC (Blau et al., 2010; Blau, van Atteveldt, Ekkebus, Goebel, & Blomert, 2009; Holloway, van Atteveldt, Blomert, & Ansari, 2015; van Atteveldt, Formisano, Goebel, & Blomert, 2004; van Atteveldt et al., 2007). In previous studies, it was named “congruency effect” when the activation in the congruent condition is stronger than the incongruent condition or “incongruency effect” when the opposite pattern was observed (Plewko et al., 2018; Wang, Karipidis, Pleisch, Fraga-Gonzalez, & Brem, 2020). To avoid confusion, we used the term “neural integration” and presented the direction if applicable. Compared with additive enhancement, neural integration appears later and is observed only when the information from different modalities is successfully integrated (Xu et al., 2019). Therefore, it reflects deep processes on the information carried by the stimuli. For example, although the involvement of STC in audiovisual integration of print and speech sound has been replicated regardless of the level of processing (e.g., phoneme, syllable, or whole word) (Kronschnabel et al., 2014; McNorgan et al., 2013; Wang et al., 2020), the precise manifestation of the effect (e.g., direction, strength) and whether other areas display such an effect are associated with multiple factors, such as orthographic depth (Holloway et al., 2015; Xu, Kolozsvari, Monto, & Hämäläinen, 2018) and reading ability (Blau et al., 2010; Blau et al., 2009; Kronschnabel et al., 2014), which are of interest in the current study. In addition, task paradigm (McNorgan & Booth, 2015; van Atteveldt et al., 2007), stimuli properties (Kronschnabel et al., 2014), and developmental stage (Plewko et al., 2018; Wang et al., 2020) also affect neural integration but are beyond the scope of this study.

Orthographic depth describes the consistency of the mapping between orthographic and phonological codes. It significantly influences behavioral and neural profiles of reading development and impairments in alphabetic languages (Borleffs, Maassen, Lyytinen, & Zwarts, 2018; Richlan, 2014). The neural audiovisual integration is also modulated by orthographic transparency (van Atteveldt & Ansari, 2014). For example, native speakers of English (deep) show a significant neural integration in the “congruent < incongruent” direction in the STC during processing letter-sound pairs, which is different from the neural integration in the “congruent > incongruent” direction observed in Dutch (shallow) (Holloway et al., 2015). More importantly, when the same group of English-speaking participants processed numbers and number names, where the mapping was one-on-one, the “congruent > incongruent” pattern appeared. Based on previous findings in alphabetic languages with varying orthographic transparency (van Atteveldt & Ansari, 2014), a neural integration in the “congruent < incongruent” direction during character-sound integration was expected. Furthermore, since semantic information is involved in character recognition, we expected additional regions such as the inferior and middle frontal cortices involved in semantic processing (Wu, Ho, & Chen, 2012; Zhao et al., 2014) to be recruited. In contrast, pinyin-sound integration was expected to induce neural integration in the “congruent > incongruent” direction.

In addition to investigating the neural basis of print-sound integration at the group level, understanding how it correlates to individual heterogeneity in reading abilities is valuable theoretically and practically. One primary reason is that skill-dependent processing is often masked by averaging at the group level (Kanai & Rees, 2011; Kidd, Donnelly, & Christiansen; Seghier & Price, 2018). Such information can also help clarify the functional significance of the observed effect. In terms of reading abilities, large individual differences are noted and can be accounted for by variances at the brain level (Kast, Bezzola, Jancke, & Meyer, 2011; McNorgan, Awati, Desroches, & Booth, 2014; McNorgan et al., 2013; Plewko et al., 2018; Wang et al., 2020). Nowadays, while a large body of research on neural correlates of print-sound integration compared children with and without RD and revealed a reduced neural integration in the RD group (e.g., Blau et al., 2010; Blau et al., 2009; Plewko et al., 2018), the relationships between the neural integration of print-sound and reading abilities within typically developing children starve for further investigations. Therefore, we additionally adopted the individual differences approach to investigate whether and how do neural circuitries link the neural integration in processing character/pinyin-sound pairs with typical reading acquisition.

## Methods

### Participants

The initial sample consisted of 54 typically developing children according to the following inclusion criteria: (a) native speaker of Mandarin Chinese, (b) right-handed, (c) normal audition, (d) normal or corrected-to-normal vision, (e) full-scale intelligence quotient (IQ) ≥ 80, (f) no history of neurological or psychiatric disorders, and (g) no reading difficulties (> 16^th^ percentile on the standardized reading screening task *Character Recognition*; i.e., above -1 standard deviation [*SD*] of the norm) (Xue, Shu, Li, Li, & Tian, 2013). All the children received formal instruction on pinyin in the first half of the semester in the 1^st^ grade. Before the experiment, the children and their parents/guardians were informed about the aim and procedure in detail and signed written consent. All behavioral and neuroimaging data were collected during 2018. This study received ethical approval from the Institutional Review Board of State Key Laboratory of Cognitive Neuroscience and Learning at Beijing Normal University.

The quality of each functional run was evaluated based on Children’s performance in the in-scanner task and the number of outliers (i.e., “bad data points”) in blood-oxygen-level-dependent signals caused by severe head motion or spikes. Acceptance depended on the following criteria: (a) no more than 15% of the volumes (10 data points) were marked as outliers; (b) no less than 75% in overall accuracy (combining all the conditions) and accuracy for each condition should be no less than 50%. The run failing to meet either criterion was removed from the analysis. As a result, 36 children had acceptable imaging data for the character experiment (23 girls; age 110-141 months; mean age = 127, grades 3-5). Eighteen children were excluded due to incomplete data collection (n = 9), poor data quality (n = 8), or the combination of poor data quality and task performance (n = 1). The final sample in the pinyin experiment consisted of 41 children (27 girls; age 110-141 months; mean age = 126, grades 3-5). Thirteen children were excluded due to incomplete data collection (n = 6), poor data quality (n = 6), or the combination of poor data quality and task performance (n = 1).

### Behavior measures

Each child received a battery of behavior tests individually in a silent booth. IQ was measured with the abbreviated version of the *Chinese Wechsler Intelligence Scale for Children* (*WISC-CR*) (Wechsler, 1974). Specifically, *Information, Similarities*, and *Digit Span* subtests were used to estimate verbal IQ; *Picture Completion, Block Design*, and *Coding* subtests were used to estimate performance IQ.

A set of neuropsychological tasks was used to measure children’s reading and reading-related cognitive-linguistic skills. In brief, *Character Recognition* used to identify Chinese dyslexic children (Cui, Xia, Su, Shu, & Gong, 2016; Xia, Hoeft, Zhang, & Shu, 2016) was used for screening aim. *Word List Reading* and *Silent Reading Comprehension* measured oral reading fluency and reading comprehension proficiency, respectively. The former relies more on grapho-phonological mapping, whereas the latter relies more on grapho-semantic processing (Xia et al., 2018). Finally, *Phoneme Deletion, Rapid Naming (RAN)*, and *Morphological Production* tasks were used to measure phonological awareness (PA), RAN, and morphological awareness (MA), the three critical cognitive-linguistic skills in Chinese reading development (Lei et al., 2011).

*Character Recognition* is a standardized test for estimating the number of characters children have learned. The test consisted of 150 characters selected from grades 1-6 textbooks (Xue et al., 2013). The characters were arranged in the order of difficulty on ten sheets of A4 paper. The participant was asked to name the characters in sequence until failing in all 15 items on one page during the task. There was no time limit. Each correct answer is worth 1 point, and the full mark is 150.

*Word List Reading* is a standardized test for measuring oral reading fluency (Zhang et al., 2012). The test consisted of 180 high-frequency two-character words arranged in a 9-column by 20-row matrix in a sheet of A4 paper. The participant was required to read out the words in left-to-right, up-to-down order as accurately and quickly as possible. The time used to complete the task was recorded, and an index referring to the number of words the participant read correctly per minute was calculated.

*Silent Reading Comprehension* is a standardized test for assessing children’s reading comprehension (Lei et al., 2011). The test included 100 sentences or short passages arranged based on the number of characters. The participant was requested to read each item silently and decide the correctness of the meaning with a mark of ✔ or ✗ as fast as possible within 3 minutes. The total number of characters in sentences with correct responses was calculated and then transformed to the number of characters the participant read per minute.

*Phoneme Deletion* was used to assess PA (H. Li, Shu, McBride-Chang, Liu, & Peng, 2012). This task included three parts, and the participant was asked to delete the initial, middle, or final sound from the orally presented syllable and pronounce the remaining portion. Each part consisted of 3 practice trials and 8-10 testing items. One correct response is worth 1 point for a total possible score of 28.

*Digit RAN* required the participant to name 50 one-digit numbers on a card as accurately and rapidly as possible (Liu et al., 2017). Numbers 1, 2, 3, 5, and 8 were repeated ten times each and arranged randomly in a 5-column by 10-row matrix. The task was administered twice, and the average time the participant used to complete the task was calculated as the final score.

*Morphological Production* was used to measure MA at the character level (Shu, McBride-Chang, Wu, & Liu, 2006). In each trial, a character was orally presented to the participant in a high-frequency word. Then, the participant had to respond with two new words. In one word, the given character retains the same meaning as in the original case, whereas in the other word, the meaning of the character should be different. This task contained 15 characters for a total possible score of 30.

### Stimuli and experimental design

In this study, we investigated the neurofunctional correlates of audiovisual integration in processing character-sound and pinyin-sound pairs. In the character experiment, the materials were characters and corresponding speech sounds (syllables) (**Figure 1A**). Fifty-six high frequency pictographic characters were selected (Chinese Single-Character Word Database; https://blclab.org/pyscholinguistic-norms-database/). All the characters are visually simple, taught early, and used as radicals in compound characters. During the experiment, visual stimuli were displayed in white, “KaiTi” font, and 96 pt at the center of a black background. A native Chinese male recorded the speech sounds with a sampling rate of 44.1 kHz and 16-bit quantization. The audio files were normalized to 85 dB and then edited with a bandpass (100-4000 Hz) filter with Audacity (https://www.audacityteam.org/). The average duration of all sounds was 476.3 (±87.5) ms without pre- or post-silence. All the speech sounds can be recognized easily. In the pinyin experiment, the visual stimuli were syllables written in pinyin (e.g., sound /sheng 1/ can be written as “shēng”, **Figure 1B**), in white, “Century Schoolbook” font, and 90 pt. The auditory stimuli were the same as those used in character conditions.

**Figure 1.**
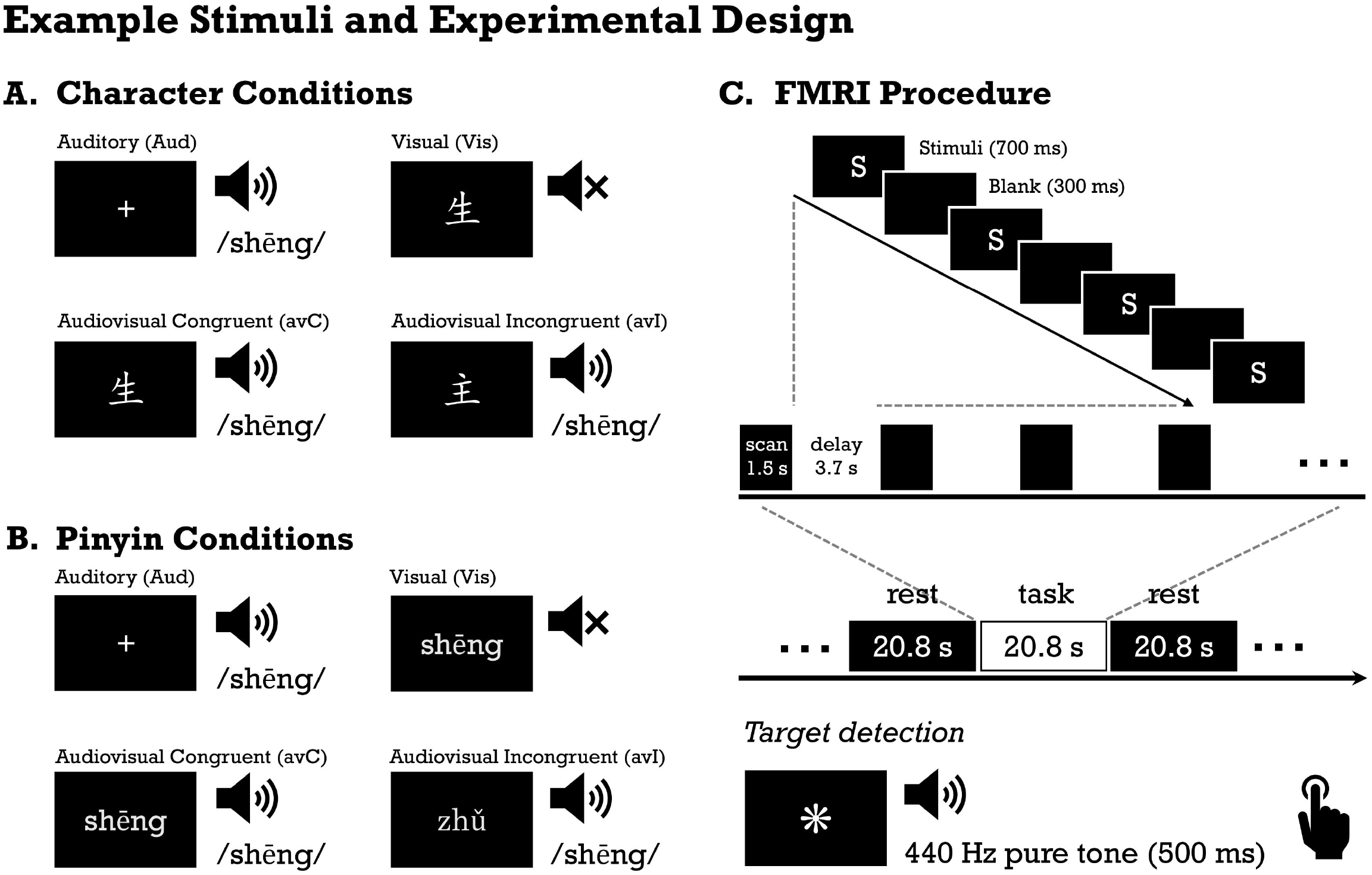
Materials and experimental design. (A) Example stimuli in the character conditions. (B) Example stimuli in the pinyin conditions. (C) Schematic illustration of the fMRI procedure and auditory and visual targets in the in-scanner task.

The fMRI procedure (**Figure 1C**) was adapted from Blau et al. (2010). A block design was used with four conditions in each experiment: unimodal auditory (Aud), unimodal visual (Vis), audiovisual congruent (avC), and audiovisual incongruent (avI). The visual and auditory stimuli were presented simultaneously in the avC and avI conditions. The entire study contained 4 functional runs, half of which were for character and half for pinyin. To avoid priming from character stimuli on pinyin processing, the pinyin experiment was always conducted first. Each run included 8 experimental blocks (duration 20.8 s), with each condition replicated twice and interleaved with 9 rest blocks (duration 20.8 s). In each experimental block, there were 4 mini-blocks. Each mini-block consisted of a 1.5-s brain volume acquisition period and a 3.7-s silence period (i.e., sparse-sampling), during which 4 trials of stimuli were presented. The order of the stimuli and conditions were pseudorandomized. Considering the age of the participants, a target detection task that does not require active monitoring of congruency status was used to help maintain attention during scanning. Compared to the active matching task, the passive paradigm will not change the automatic response pattern during audiovisual integration (Blau et al., 2010; van Atteveldt et al., 2007). The participant was asked to press the button with the right index finger accurately and quickly to the auditory (440 Hz pure tone), visual (an unpronounceable symbol) targets, or their combinations that appeared twice randomly per experimental block. Two practice sessions were administered to ensure that the child performed the task correctly, including one outside the scanner and the other before the first fMRI session inside the scanner.

### Image acquisition

The 3-Tesla Siemens MAGNETOM Trio Tim scanner in Imaging Center for Brain Research at Beijing Normal University was used to collect all the images with a 12-channel head coil. For each child, 4 T2^*^ sensitive gradient echo planar imaging sequences were collected with the following parameters: repetition time (TR) = 5200 ms (consist of 1500 ms for image acquisition and 3700 ms delay), echo time = 32 ms, flip angle = 90 degrees, slice thickness = 4.5 mm, voxel size = 3.0 × 3.0 × 4.5 mm^3^, interscan gap = 0.675 mm, number of slices = 24, number of volumes = 68, and time of acquisition = 5 minutes and 54 s. The sparse-sampling method provided a 3.7-s period of silence for stimuli presentation. Such a design is recommended for fMRI experiments investigating processing with auditory inputs (Talavage & Hall, 2012). Three dummy scans were administered before image collection for each functional run to avoid scanner equilibration effects. In addition, a high-resolution whole-brain T1-weighted structural image (magnetization-prepared rapid acquisition with gradient echo, TR = 2530 ms, echo time = 3.39 ms, inversion time = 1100 ms, flip angle = 7 degrees, slice thickness = 1.33 mm, voxel size = 1.33 × 1.0 × 1.33 mm^3^, number of axial slices = 144, and time of acquisition = 8 minutes and 7 s) was collected. Prior to the formal scan, children were informed about the experimental procedure, familiarized with the noise of MRI sequences, and trained to hold still. Earplugs and noise-proof headphones were used to help reduce scanner noise, and foam pads were used to help reduce head motion during scanning. A radiologist blinded to the details of the study reviewed all the images to ascertain any pathological deviations.

### FMRI data preprocessing

Preprocessing of brain images was conducted with SPM12 (Wellcome Department of Cognitive Neurology, London, UK, http://www.fil.ion.ucl). It included the following steps: (a) head motion correction (realign), (b) ART-based outlier detection (the “bad time point” was defined as intensity > global mean ± 9 SD or frame-to-frame head motion [framewise displacement] > 2 mm; https://www.nitrc.org/projects/artifact_detect), (c) T1 image segmentation, (d) normalization to the standard template in Montreal Neurological Institute (MNI) space, (e) smoothing with an 8-mm full-width at half-maximum Gaussian kernel. No slice timing correction was performed, given the discontinuous nature of the signal in fMRI data collected using sparse-sampling (Perrachione & Ghosh, 2013). At the individual level, experimental conditions were modeled with a generalized linear model. Parameters referring to outliers in the time course and head motion (3 translation, 3 rotation, and 1 framewise displacement) were included in the model to exclude the effect of nuisance covariates. Then, we calculated two brain maps— avC vs. (Aud + Vis) / 2 and avC vs. avI, which were used in the subsequent analyses on additive enhancement and neural integration, respectively.

### Statistical analyses

Descriptive statistics of demographic and behavioral measures were performed, followed by calculating correlations between the reading abilities (*Character Recognition, Word List Reading, Silent Reading Comprehension*) and cognitive-linguistic skills (*Phoneme Deletion, RAN, Morphological Production*). The Bonferroni method was used to control multiple comparisons error (corrected *p* < 0.05 / 15). Next, we evaluated in-scanner task performance. Analysis of variance (ANOVA) was performed separately on accuracy and reaction time (RT) with run and condition as repeated measures. In this study, we defined the accurate response as a button press to the target with an RT between 200 and 2000 ms. The average reaction time of correct responses was calculated.

Additive enhancement and neural integration were used to identify brain areas involved in the audiovisual processing of character and pinyin stimulations, respectively. Given that these indicators reflect different aspects of multisensory processing, combining them enables distinguishing regions with different functions. Furthermore, we examined the relationships between neural integration with reading abilities and cognitive-linguistic skills to deepen understanding of specific regions in reading processing. In brief, we first conducted voxel-wise whole-brain analyses to identify regions showing additive enhancement, neural integration, and integration-reading correlation in each experiment. Next, we performed conjunction analysis to identify areas showing single or multiple effects. These regions were then defined as regions-of-interest (ROIs), in which subsequent analyses were conducted. Details about each part are presented below.

In the voxel-wise whole-brain analysis, we first searched for regions showing mean additive enhancement with two criteria: (a) active in both unimodal conditions (i.e., Aud > 0, Vis > 0). This criterion was used to exclude areas that only responded to one input modality. (b) stronger activation in the audiovisual congruent condition than the mean of unimodal conditions, i.e., avC > (Aud + Vis) / 2. Three one-sample *t*-tests were performed, and the intersections were located. For each *t*-test, the uncorrected threshold of *p*-voxel < 0.05 was used, resulting in a conjoint *p* < 0.05^3 = 0.000125 for the overlapping voxel. In common practice, the intersection map will not be thresholded. However, to be conservative, we reported and discussed clusters containing more than 200 continuous voxels (i.e., 1,600 mm^3^; arbitrarily defined). Next, we identified regions showing neural integration (i.e., brain responses differed between the congruent condition against incongruent condition). A one-sample *t*-test was performed with individuals’ contrast maps of “congruent vs. incongruent.” We focused on brain areas showing activation (uncorrected *p*-voxel < 0.05) in at least one multimodal condition to avoid results produced by differences in de-activation. A threshold of *p*-voxel < 0.005, Family-Wise Error (FWE) corrected *p*-cluster < 0.05 was used to control multiple comparisons error. Previous studies repeatedly reported that neural integration was associated with reading ability. Thus, to avoid missing the effect masked at the group level, we conducted two regression analyses where the standard scores of oral reading fluency and silent reading comprehension proficiency were included as the variate of interest, respectively. The same threshold of FWE corrected *p* < 0.05 (cluster-forming *p*-voxel < 0.005) was used. Nuisance variables of age and sex were controlled statistically in all the analyses.

We next performed conjunction analysis with the results of whole-brain analyses to identify areas showing single or multiple effects. Such information will shed light on the role each area plays in print-sound processing. Since domain-specific (i.e., reading-related) processes were of interest in this study, we focused on areas with at least one neural integration-related effect (i.e., showing neural integration in either direction and/or significant correlation between neural integration and reading abilities) and defined them as ROIs. The primary aim of the ROI analysis was to visualize certain effects. The second aim was to examine whether the absence of significance on a specific effect was due to the strict whole brain threshold. Specifically, in the ROI that did not show additive enhancement in the whole brain analysis, we examined the average activation in each unimodal condition and re-examined the additive enhancement with the average activation within the region. Similarly, in the ROI that did not show neural integration in the whole brain analysis, we re-examined the neural integration with the average values. In the ROI that did not show any integration-reading correlation in the whole brain analyses, we re-examined the relationships at the ROI level while using the Bonferroni method to control the number of behavior variables (two reading ability and three cognitive-linguistic skills: corrected *p* < 0.05 / 5 = 0.01). The third aim of ROI analysis was to examine the specificity of the brain-behavior correlation revealed in the whole-brain analysis. For example, for the ROI where the neural integration was correlated oral reading fluency, we then calculated its correlation with the other reading ability (silent reading comprehension proficiency) and cognitive-linguistic skills (PA, RAN, and MA), while the nuisance variables of age and sex were controlled statistically. Bonferroni correction was used to control the number of behavior variables (one reading ability and three cognitive-linguistic skills; corrected *p* < 0.05 / 4 = 0.0125). Finally, as the left STC was identified in both the character experiment (showing additive enhancement and neural integration) and the pinyin experiment (showing additive enhancement and integration-reading correlation), we performed conjunction analysis to examine the spatial overlap between the two clusters. We also calculated the correlation between the magnitudes of neural integration in processing character-sounds and pinyin-sounds. Of note, in the ROI that was defined by a certain effect, we would not re-examine this effect. Plots were created only for the visualization aim.

Significant clusters identified in the whole brain analyses were presented over a FreeSurfer surface template with BrainNet Viewer (Xia, Wang, & He, 2013). The peak location was reported in MNI coordinates, and corresponding brain regions were localized and reported with the AAL atlas implemented in DPABI (http://rfmri.org/dpabi). Behavioral analysis and ROI analysis were performed with SPSS Statistics version 24 (SPSS Inc., Chicago, IL, USA).

## Results

### Behavioral profiles and in-scanner task performance

Descriptive statistics of demographics, cognitive-linguistic and reading skills are presented in **Table 1**. Significantly positive correlations were observed between reading measures (all *p*’s < 0.05 / 15, Bonferroni correction; see **Table 2** for the correlation coefficients and raw *p*-values). Regarding the relationship with cognitive-linguistic skills, *Word List Readin*g was significantly associated with *Digit RAN* (*r* = -0.597, *p* < 0.001 < 0.05 / 15), but not with *Phoneme Deletion* (*r* = 0.288, *p* = 0.068 > 0.05 / 15) or *Morphological Production* (*r* = 0.344, *p* = 0.028 > 0.05 / 15) with Bonferroni correction. In contrast, *Silent Reading Comprehension* was significantly correlated with morphological production (*r* = 0.487, *p* = 0.001 < 0.05 / 15), but not with *Phoneme Deletion* (*r* = 0.135, *p* = 0.400 > 0.05 / 15) or *Digit RAN* (*r* = -0.332, *p* = 0.034 > 0.05 / 15) with Bonferroni correction. This pattern indicates that in typically developing Chinese children who have received formal reading instruction for 3-5 years, oral reading relies more on print-to-sound mapping, whereas reading comprehension relies more on semantic processing.

**Table 1.** Demographics and behavior profiles (n = 41)

| Measure | *Mean* | *SD* | *Range* |
| --- | --- | --- | --- |
| Age (month) | 126 | 8 | 110-141 |
| Grade (3<sup>rd</sup> / 4<sup>th</sup> / 5<sup>th</sup>) |  | 4 / 23 / 14 |  |
| Sex (female / male) |  | 27 / 14 |  |
| Verbal IQ (standard score) | 107 | 13 | 85-145 |
| Performance IQ (standard score) | 114 | 13 | 75-137 |
| Full-scale IQ (standard score) | 111 | 12 | 80-143 |
| Character Recognition (corrected item) | 119 | 12 | 86-146 |
| Word List Reading (word / minute) | 90 | 19 | 55-140 |
| Silent Reading Comprehension (character / minute) | 327 | 119 | 132-565 |
| Phoneme Deletion (corrected item) | 21 | 4 | 7-25 |
| Rapid Naming (second) | 18 | 3 | 11-24 |
| Morphological Production (item) | 24 | 4 | 13-30 |
| Number of “bad time points” (outliers) per fMRI run |  |  |  |
| Character Experiment <sup>a</sup> | 2.4 | 3.37 | 0-10 |
| Pinyin Experiment | 0.74 | 1.54 | 0-9 |

*Note.* <sup>a</sup> n = 36

**Table 2.** Correlation between reading and cognitive-linguistic skills (n = 41)

| Measures | 1 | 2 | 3 | 4 | 5 | 6 |
| --- | --- | --- | --- | --- | --- | --- |
| 1 Character Recognition | --- |  |  |  |  |  |
| 2 Word List Reading | $r = 0.559$<br>$p < 0.001$ | --- | | | | |
| 3 Silent Reading Comprehension | $r = 0.647$<br>$p < 0.001$ | $r = 0.701$<br>$p < 0.001$ | --- | | | |
| 4 Phoneme Deletion | $r = 0.286$<br>$p = 0.070$ | $r = 0.288$<br>$p = 0.068$ | $r = 0.135$<br>$p = 0.400$ | --- | | |
| 5 Rapid Naming | $r = -0.357$<br>$p = 0.022$ | $r = -0.597$<br>$p < 0.001$ | $r = -0.332$<br>$p = 0.034$ | $r = -0.160$<br>$p = 0.319$ | --- | |
| 6 Morphological Production | $r = 0.566$<br>$p < 0.001$ | $r = 0.344$<br>$p = 0.028$ | $r = 0.487$<br>$p = 0.001$ | $r = 0.122$<br>$p = 0.446$ | $r = -0.117$<br>$p = 0.466$ | --- |

All the children performed well in the in-scanner task (character accuracy: *Mean* [*SD*] = 96.5% [4.8], RT: *Mean* [*SD*] = 512 [81] ms; pinyin accuracy: *Mean* [*SD*] = 96.4% [3.7], RT: *Mean* [*SD*] = 530 [80] ms; **Table S1**). No significant differences were found between runs (all *p*’s > 0.1). The results of ANOVA with condition as the repeated measure factor revealed no significant effect on accuracy (*F* = 1.00, *p* = 0.395) and a significant effect on RT in the character experiment (*F* = 35.8, *p* < 0.001). Post-hoc tests revealed the longest RT for the unimodal visual condition (*p*’s ≤ 0.002 in all comparisons, Bonferroni correction), followed by the auditory condition (Aud > avC: corrected *p* < 0.001; Aud > avI: corrected *p* < 0.001). There was no difference between the two crossmodal conditions. In the pinyin experiment, the effect of condition was significant on both accuracy (*F* = 10.1, *p* < 0.001) and RT (*F* = 54.1, *p* < 0.001). Post-hoc tests revealed that while children had the lowest accuracy in the visual condition (*p*’s ≤ 0.005 in all comparison, Bonferroni correction), no differences were found between any other pairs of conditions. In the unimodal visual condition, participants also showed the longest RT (all corrected *p*’s ≤ 0.05), followed by the auditory condition (Aud > avC: corrected *p* < 0.001; Aud > avI: corrected *p* < 0.001). No differences were observed between the two crossmodal conditions.

### Brain results: the character experiment

The bilateral STCs were identified showing additive enhancement (**Figure 2A**; **Table 3**; see **Figure S1** for activation in two unimodal conditions and the brain map for avC > (Aud + Vis) / 2). On the other hand, a significantly neural integration (avC < avI) was found in the left frontal (peak MNI -40, 14, 32), parietal (peak MNI -36, -62, 48) and bilateral STCs (peak MNI, left: -54, -22, -2; right: 62, -34, 10; **Figure 2B, Table 4**) at the FWE corrected threshold of *p*-cluster < 0.05. In addition, whole-brain regression analysis identified a cluster in the left middle frontal cortex (peak MNI -32, 10, 42), showing a positive correlation between the neural integration and silent reading comprehension proficiency at the FWE corrected threshold of *p*-cluster < 0.05 (**Figure 2C, Table 4**). No regions were identified as significant in whole-brain regression with oral reading fluency as the variable of interest.

**Table 3.** Cluster identified showing audiovisual additive enhancement in each experiment

| Experiment | Number of Voxels | Brain area <sup>a</sup> |
| --- | --- | --- |
| Character | 822 | Temporal_Sup_R, Temporal_Mid_R |
|  | 354 | Temporal_Sup_L, Temporal_Mid_L |
| Pinyin | 2752 | Temporal_Mid_L, Temporal_Sup_L, Rolandic_Oper_L, Occipital_Mid_L, Temporal_Inf_L, SupraMarginal_L, Fusiform_L, Postcentral_L, Heschl_L, Temporal_Pole_Sup_L, Occipital_Inf_L |
|  | 1532 | Temporal_Sup_R, Temporal_Mid_R, SupraMarginal_R |
|  | 967 | Temporal_Mid_R, Temporal_Inf_R, Fusiform_R, Cerebelum_6_R, Occipital_Inf_R, Occipital_Mid_R |
|  | 943 | Precentral_L, Frontal_Inf_Oper_L, Frontal_Inf_Tri_L, Rolandic_Oper_L |
|  | 936 | Putamen_L, Thalamus_L, Insula_L, Vermis_3 |

*Note.* <sup>a</sup> labeling is based on the AAL atlas (> 20 voxels).

**Table 4.**
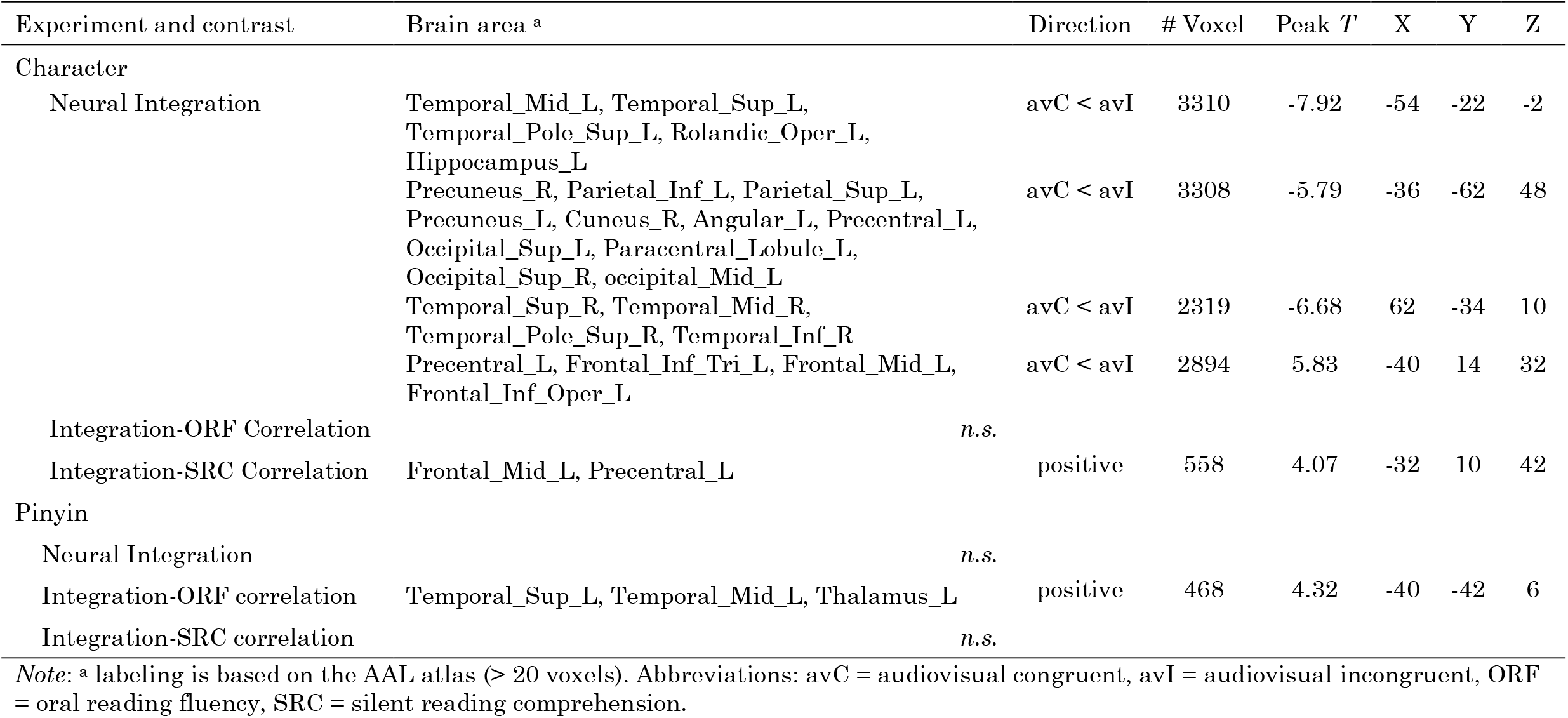
Significant clusters identified in the voxel-wise whole-brain analyses on neural integration and integration-reading correlation

| Experiment and contrast | Brain area <sup>a</sup> | Direction | # Voxel | Peak *T* | X | Y | Z |
| --- | --- | --- | --- | --- | --- | --- | --- |
| Character |  |  |  |  |  |  |  |
| Neural Integration | Temporal_Mid_L, Temporal_Sup_L, Temporal_Pole_Sup_L, Rolandic_Oper_L, Hippocampus_L | avC < avI | 3310 | -7.92 | -54 | -22 | -2 |
|  | Precuneus_R, Parietal_Inf_L, Parietal_Sup_L, Precuneus_L, Cuneus_R, Angular_L, Precentral_L, Occipital_Sup_L, Paracentral_Lobule_L, Occipital_Sup_R, occipital_Mid_L | avC < avI | 3308 | -5.79 | -36 | -62 | 48 |
|  | Temporal_Sup_R, Temporal_Mid_R, Temporal_Pole_Sup_R, Temporal_Inf_R | avC < avI | 2319 | -6.68 | 62 | -34 | 10 |
|  | Precentral_L, Frontal_Inf_Tri_L, Frontal_Mid_L, Frontal_Inf_Oper_L | avC < avI | 2894 | 5.83 | -40 | 14 | 32 |
| Integration-ORF Correlation |  | *n.s.* |  |  |  |  |  |
| Integration-SRC Correlation | Frontal_Mid_L, Precentral_L | positive | 558 | 4.07 | -32 | 10 | 42 |
| Pinyin |  |  |  |  |  |  |  |
| Neural Integration |  | *n.s.* |  |  |  |  |  |
| Integration-ORF correlation | Temporal_Sup_L, Temporal_Mid_L, Thalamus_L | positive | 468 | 4.32 | -40 | -42 | 6 |
| Integration-SRC correlation |  | *n.s.* |  |  |  |  |  |

*Note:* <sup>a</sup> labeling is based on the AAL atlas (> 20 voxels). Abbreviations: avC = audiovisual congruent, avI = audiovisual incongruent, ORF = oral reading fluency, SRC = silent reading comprehension.

**Figure 2.**
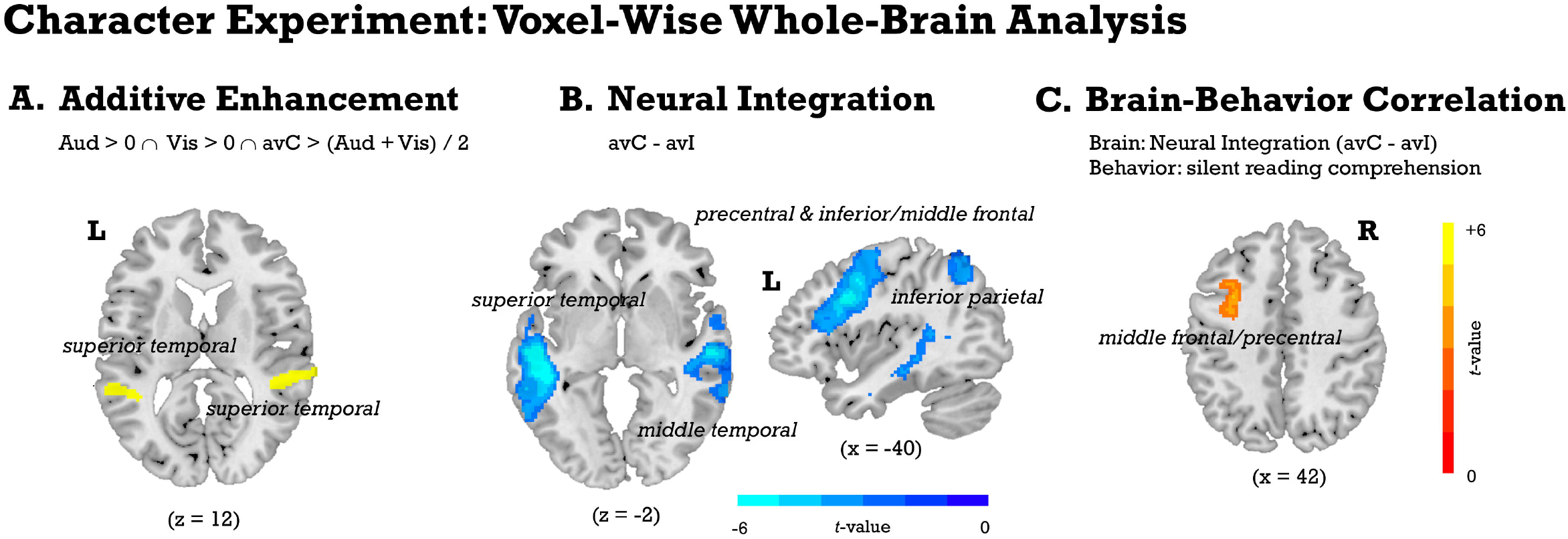
The whole-brain results of the character experiment. (A) Brain areas showing audiovisual additive enhancement are identified with the mean rule (Beauchamp, 2005): (1) activate in both unisensory conditions (i.e., Aud > 0, Vis > 0); (2) audiovisual congruent stimuli induce stronger activation than the average activation of the auditory and visual conditions (i.e., avC > (Aud + Vis) / 2). The uncorrected threshold of *p*-voxel < 0.05 is used for each contrast map, resulting in a conjoint probability of *p*-voxel < 0.05^3 = 0.000125. The cluster containing no less than 200 continuous voxels under the threshold is considered to show additive enhancement and is colored yellow. (B) Brain areas showing a significant neural integration (blue represents the “avC < avI” direction). Threshold: cluster forming *p*-voxel < 0.005, Family-Wise Error (FWE) corrected *p*-cluster < 0.05. No regions were found displaying a significant neural integration in the “avC > avI” direction. (C) The brain area shows a significant correlation between neural integration (avC - avI) and silent reading comprehension proficiency. Threshold: cluster forming *p*-voxel < 0.005, FWE corrected *p*-cluster < 0.05. *Abbreviations*: Aud = auditory, Vis = visual, avC = audiovisual congruent, avI = audiovisual incongruent, L = left hemisphere, R = right hemisphere.

Based on results produced by the whole brain analyses, we defined three types of regions. The first type (bilateral STCs) showed both additive enhancement and neural integration (**Figure 3A-B**). No correlations with reading scores were found even at uncorrected *p* < 0.05 in the ROI analysis. The second type of region (left inferior parietal lobule) showed only neural integration in the whole brain analyses (**Figure 3C**). We found no activation in either the unimodal condition or the audiovisual congruent condition. No brain-behavior correlations were found in this area at the ROI level neither. The third type of region (left middle frontal gyrus; MFG) showed a significant neural integration, of which the magnitude was associated with silent reading comprehension proficiency (**Figure 3D**). The ROI analysis found significant activation for unimodal auditory and visual conditions in this area but observed no activation in the audiovisual congruent condition. For the brain-behavior relationship, there was no significant correlation under the FWE correction. The magnitude of neural integration also correlated with oral reading fluency (*r* = 0.382, *p* = 0.026) at an uncorrected threshold, and showed correlated trends for digit RAN (*r* = -0.319, *p* = 0.066) and MA (*r* = 0.300, *p* = 0.084). Importantly, the positive correlation between neural integration and silent reading comprehension remained significant (*r* = 0.494, *p* = 0.004) even after controlling for oral reading fluency. Finally, we found that the positive correlation between neural integration and reading comprehension was driven by a negative correlation in the incongruent condition (avC: *r* = 0.084, *p* = 0.637; avI: *r* = -0.487, *p* = 0.003).

**Figure 3.**
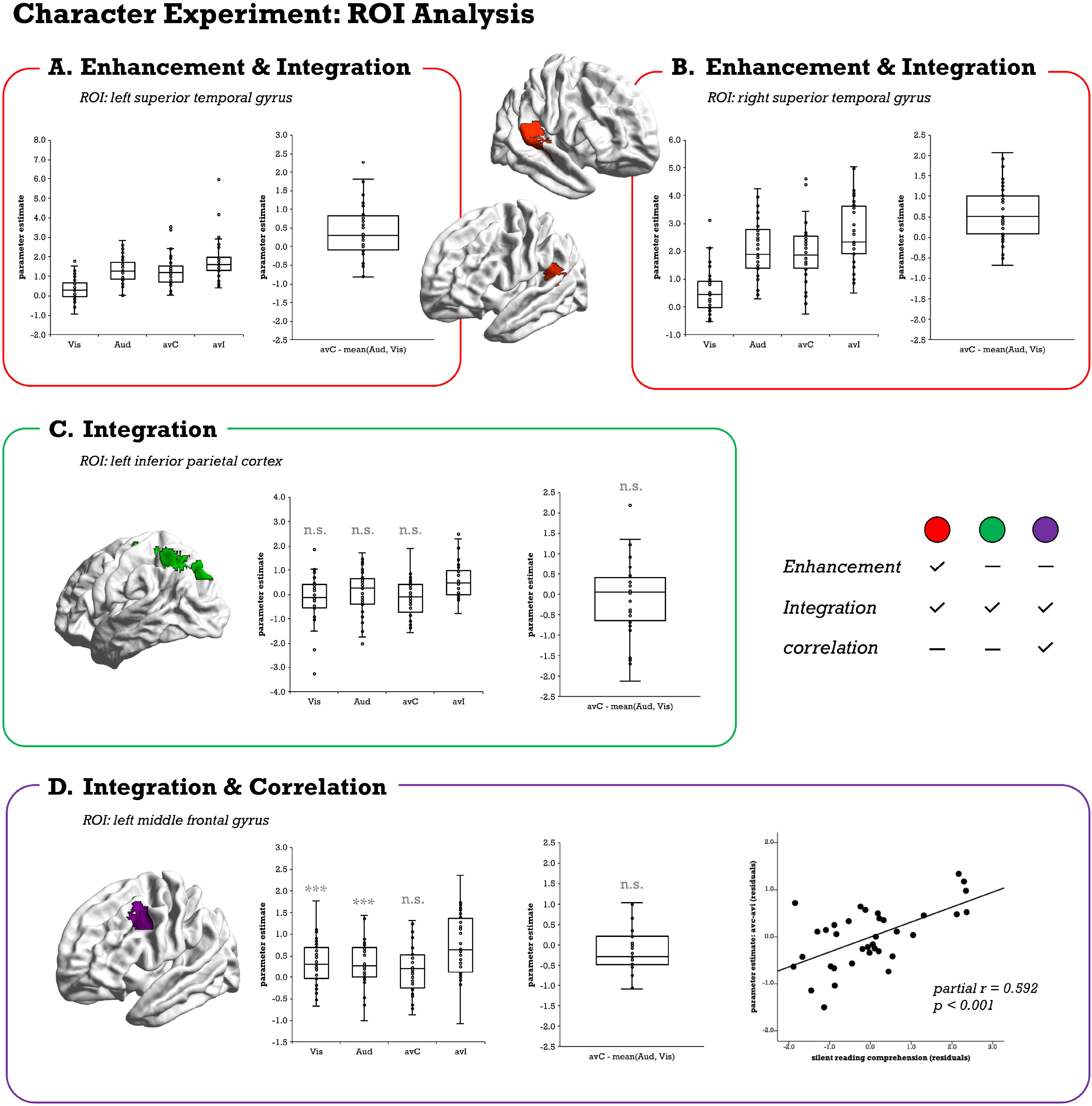
Region-of-interest (ROI) analyses of the character experiment. The intersection between brain areas showing additive enhancement and neural integration (A and B), neural integration only (C), and both neural integration and integration-reading correlation (D) were defined as the ROIs. For each ROI, a bar plot displaying activation for each condition and a boxplot displaying the contrast “avC - (Aud + Vis) / 2” are presented. A scatter plot is presented in the ROI showing a significant correlation between the neural nitration and silent reading comprehension (residuals were obtained by regressing out nuisance variables of age and sex with a general linear model). *Abbreviations*: Aud = unisensory auditory, Vis = unisensory visual, avC = audiovisual congruent, avI = audiovisual incongruent; *** *p* < 0.001, *n*.*s*. = non-significant.

### Brain results: the pinyin experiment

The audiovisual additive enhancement was detected in the left superior temporal, inferior frontal, right superior, inferior temporal, and subcortical cortices (**Figure 4A**; **Table 3**; see **Figure S1** for activation in the unimodal conditions and brain map for avC > (Aud + Vis) / 2). No regions showed a significant neural integration in either direction in the whole brain analysis, which is unexpected. However, voxel-wise regression analysis revealed that oral reading fluency was positively correlated with the neural integration in the left superior temporal gyrus (STG; peak MNI -56 -36 14) (FWE corrected *p*-cluster < 0.05; **Figure 4B, Table 4**). No regions were identified as significant in whole-brain regression with silent reading comprehension proficiency as the variable of interest. Therefore, we defined the area (left STG) showing both additive enhancement and integration-reading correlation as ROI (**Figure 4C**). In the subsequent analyses, we did not observe the neural integration within this region even at the uncorrected threshold of *p* < 0.05.

**Figure 4.**
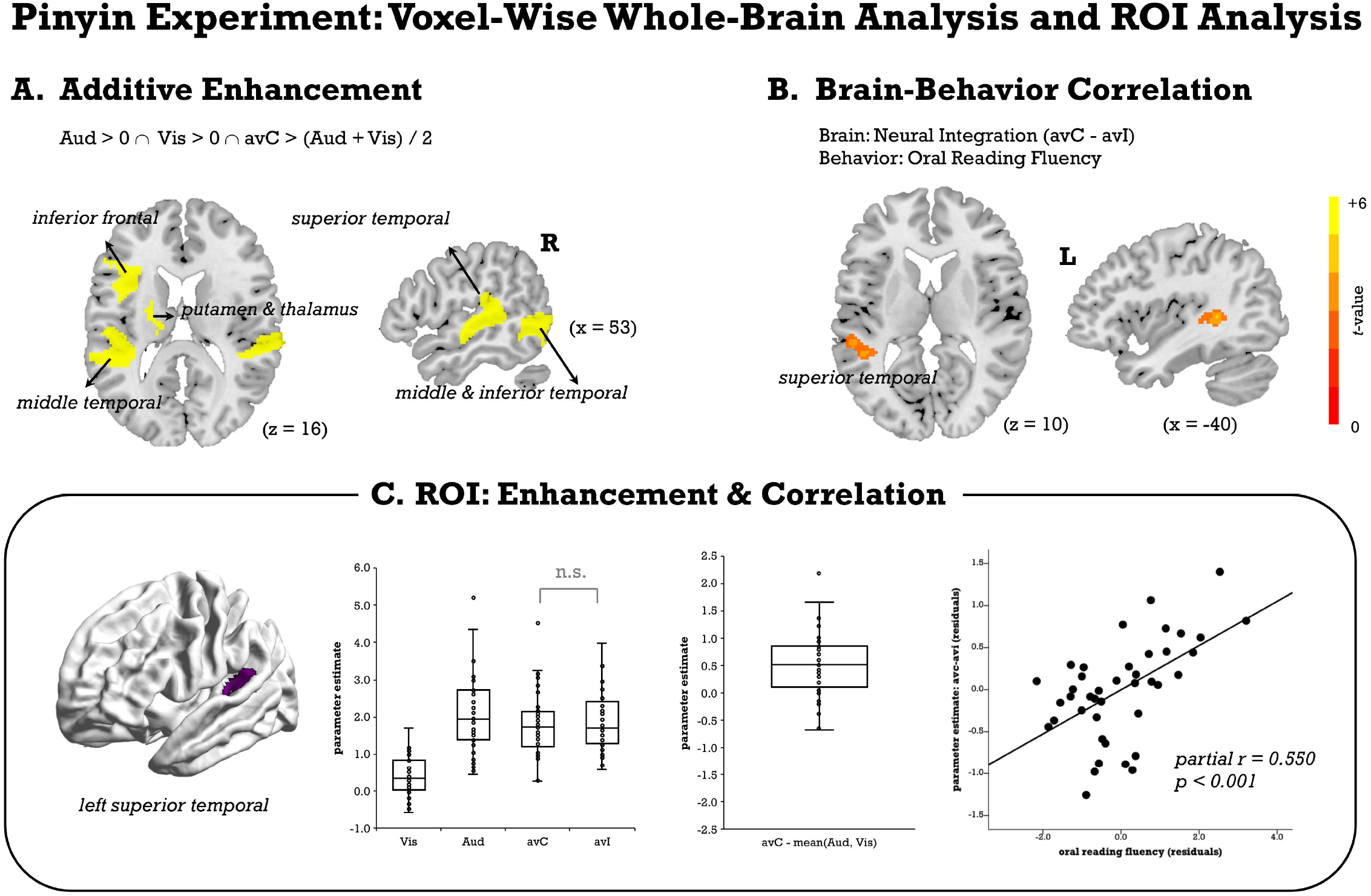
The whole-brain and region-of-interest (ROI) analyses of the pinyin experiment. (A) Brain areas showing audiovisual additive enhancement are identified with the mean rule (Beauchamp, 2005): (1) activate in both unisensory conditions (Aud > 0, Vis > 0); (2) audiovisual congruent stimuli induce stronger activation than the average activation of the auditory and visual conditions (avC > (Aud + Vis) / 2). The uncorrected threshold *p*-voxel < 0.05 is used for each contrast map, resulting in a conjoint probability of *p*-voxel < 0.05^3 = 0.000125. The cluster consisting of no less than 200 continuous voxels under the threshold is considered to show additive enhancement and is colored yellow. (B) The brain area displaying a significant correlation between neural integration (avC - avI) and oral reading fluency. Threshold: cluster forming *p*-voxel < 0.005, Family-Wise Error (FWE) corrected *p*-cluster < 0.05. (C) The intersection between areas showing additive enhancement and significant integration-reading correlation is defined as the region-of-interest (ROI). The bar plot represents activation in each condition. Boxplot displays the contrast “avC - (Aud, Vis) / 2”. The scatter plot represents the correlation between neural integration and oral reading fluency (residuals were obtained by regressing out nuisance variables of age and sex with a general linear model). *Abbreviations*: Aud = unisensory auditory, Vis = unisensory visual, avC = audiovisual congruent, avI = audiovisual incongruent, L = left hemisphere, R = right hemisphere; *n*.*s*. = non-significant.

In terms of brain-behavior correlation, no significant correlation was found after correcting for multiple comparisons error. However, the neural integration was correlated with silent reading comprehension without multiple comparisons correction (*r* = 0.371, *p* = 0.020, uncorrected). No correlations or correlated trends were found with PA (*r* = -0.087, *p* = 0.600), digit RAN (*r* = -0.248, *p* = 0.127), or MA (*r* = 0.195, *p* = 0.235). Of importance, the correlation between the neural integration and oral reading fluency remained significant (*r* = 0.437, *p* = 0.006) after additionally controlling for reading comprehension proficiency. Finally, different from the left MFG observed in the character experiment, the positive integration-reading correlation in the left STC was driven by a positive correlation in the congruent condition (avC: *r* = 0.381, *p* = 0.017; avI: *r* = -0.039, *p* = 0.815).

Given that the left STC was identified as functionally important (significant in both additive enhancement and neural integration-related analyses) in print-sound processing in the character and pinyin experiments, we conducted follow-up conjunction analysis and correlation analysis to examine their relationship. First, the two ROIs spatially overlapped (**Figure 5A)**. Second, the magnitudes of the neural integration in processing character-sound pairs and pinyin-sound pairs were significantly correlated (*r* = 0.512, *p* = 0.002; **Figure 5B**), while nuisance variables of age and sex were statistically controlled.

**Figure 5.**
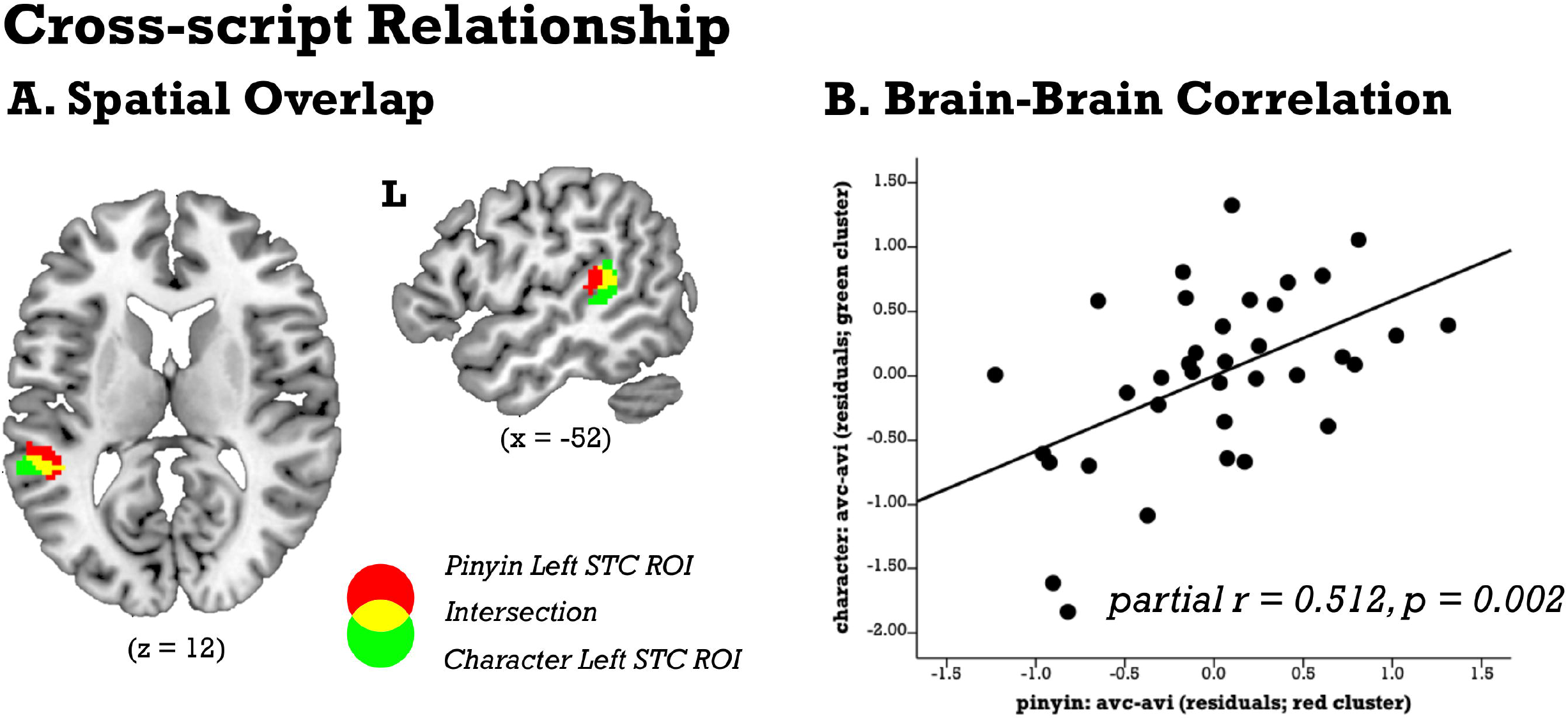
(A) The regions-of-interest (ROIs) in the left superior temporal cortex defined in the character experiment (green) and pinyin experiment (red), as well as their intersection (yellow), are presented on a template brain. (B) The scatter plot shows the positive correlation between the neural integration (avC - avI) in the character ROI and that in the pinyin ROI (residuals were obtained by regressing out nuisance variables of age and sex with a general linear model).

## Discussion

This study investigated the brain bases underlying print-sound integration in two scripts with different orthographic transparency in typically developing Chinese children. The results revealed both script-shared and script-unique neural substrates, supporting a universal principle underlying the neural mechanism of reading-related processes. While the bilateral STCs displayed additive enhancement in processing both character and pinyin categories, the differentiation between scripts was mainly revealed on the neural integration—activation differences between the congruent and incongruent conditions. Specifically, we found that the neural audiovisual integration of character-sounds in the left MFG was associated with silent reading comprehension proficiency, which might be due to the essential role of the left MFG in semantic processing and represents a language-specific manifestation in Chinese. On the other hand, the left STG links the audiovisual integration of pinyin-sounds with oral reading fluency by underpinning the shared grapho-phonological mapping component—providing a possible mechanism with which pinyin helps with Chinese reading development. This idea is further supported by the spatial overlap between the neural correlates of character-sound integration and pinyin-sound integration in the left STC and the strong correlation between the neural integration effects of the two scripts.

### Commonality and differentiation between character-sounds and pinyin-sounds processing in bilateral superior temporal cortices

This study first observed additive enhancement in the bilateral STCs in processing character-sound (opaque) and pinyin-sound (transparent) associations, suggesting a shared neural mechanism. To date, studies have investigated the neural mechanism underlying print-sound integration and its role in reading with multiple neuroimaging techniques. On the one hand, electroencephalogram and MEG are widely used to reveal the temporal properties of neural activities (Froyen, van Atteveldt, & Blomert, 2010; Froyen, Van Atteveldt, Bonte, & Blomert, 2008; Froyen, Bonte, van Atteveldt, & Blomert, 2009; Raij, Uutela, & Hari, 2000; Xu et al., 2019). These studies demonstrate that the integration occurs preconsciously in expert reading and is reflected in the components such as mismatch negativity. The fMRI research, on the other hand, provided more precise and detailed information about spatial localization. Like other stimuli categories (e.g., lip-speech, object-sound), processing audiovisual information of scripts induces super-additivity (or mean additive) enhancement in STC (Beauchamp, 2005; McNorgan & Booth, 2015; van Atteveldt et al., 2004). In line with the current results, this effect has been identified both in transparent (van Atteveldt et al., 2004) and opaque alphabetic orthographies (McNorgan & Booth, 2015). Altogether, these findings support the idea that additive enhancement is more likely to reflect fundamental and domain-general aspects of audiovisual processing.

Different from additive enhancement, script-related differences were observed on the neural integration, which is another widely used indicator of audiovisual integration and has been repeatedly reported in areas including the bilateral STCs (van Atteveldt et al., 2004; van Atteveldt, Formisano, Blomert, & Goebel, 2007). Existing studies suggest the neural integration reflects more domain-specific processing and is associated with several factors, such as linguistic characteristics (Holloway et al., 2015) and reading abilities (Blau et al., 2010; Blau et al., 2009; Karipidis et al., 2018). Consistency of grapheme-phoneme mapping (a.k.a. orthographic depth or transparency) is an essential feature in alphabetic languages. Differences in the direction of the neural integration and brain regions showing this effect have been reported between languages with different orthographic depths (Blomert & Froyen, 2010; Holloway et al., 2015). For example, stronger activation in the congruent condition than incongruent condition is more consistently reported in skilled readers when processing letter-sound associations in transparent scripts. In contrast, the opposite pattern is observed in more opaque scripts (Holloway et al., 2015). Chinese is a logographic language with an extremely deep orthography. Therefore, we expected to observe a neural integration in the “congruent < incongruent” direction. In support of this hypothesis, such an effect was identified in four regions (including the bilateral STC) in the character experiment. In contrast, in the absence of neural integration in processing pinyin-sound pairs at the group level, we observed a positive correlation between the neural integration and oral reading fluency: Higher skill readers showed a “congruent > incongruent” pattern, whereas lower skill readers showed the opposite way. This is consistent with previous studies that revealed the lack of neural integration in dyslexia in transparent languages (Blau et al., 2010; Blau et al., 2009), as well as these investigated brain-behavior correlations at both the phoneme (Karipidis et al., 2016; Karipidis et al., 2018) and syllable levels (McNorgan et al., 2014; Wang et al., 2020). In the following sections, we discussed specific neural manifestations of character-sound and pinyin-sound integration.

### Left frontal and parietal cortex: character-related neural print-sound integration

In this study, the left MFG displayed a significant neural integration and correlated with silent reading comprehension proficiency. The neural integration in the “incongruent > congruent” direction is in line with our expectation according to the opaque orthography in the Chinese writing system. Considering this region did not show additive enhancement, the brain-behavior correlation indicates the neural integration is more likely to be downstream and reflects semantic processing. This finding reflects that opaque orthography is not the only linguistic feature that affects the neural processing of character-sound processing. Chinese characters are morpheme-syllabic, which means that semantic information is deeply involved in character recognition. Therefore, in addition to grapho-phonological mapping, semantic access is an equally or even more critical aspect in reading (Bi, Han, Weekes, & Shu, 2007; Dang, Zhang, Wang, & Yang, 2018; Liu et al., 2017; Ruan et al., 2018; Yang et al., 2013; Yang et al., 2006).

Moreover, neither grapho-semantic mapping nor phono-semantic mapping is reliable at the character level in Chinese. Given that no activation was found in the audiovisual congruent condition for characters, the neural integration observed in the left frontal cortex is more likely to reflect the conflict between different meanings accessed from the visual and auditory inputs. Interestingly, the positive correlation between neural integration in the left MFG and reading comprehension proficiency was driven by a negative correlation in the incongruent condition. In other words, individuals with better comprehension performance showed lower activation for non-matching print-sound pairs. This pattern could be caused by higher neural efficiency in relevant processing, such as implicit suppression of conflicting semantic information. It has previously been found that in the initial stage of establishing a binding between grapheme and phoneme in Swiss-German, typical readers show higher activation in incongruent conditions, and such activation decreases as automation is achieved (Plewko et al., 2018).

In previous research, the frontal areas have been reported along with a broad bilateral fronto-parietal network. Its function was proposed as task-related top-down modulation (van Atteveldt et al., 2007) or implicit domain-general conflict detection and resolution (Holloway et al., 2015). Recently, the neural integration measured with MEG was sourced to the frontal cortex in experienced Chinese adult readers (Xu et al., 2019). Here, we replicated this finding by revealing such effect in the left MFG in a developmental population in Chinese with fMRI and further demonstrated that the role of this region in the implicit audiovisual integration of characters is domain-specific and that the information automatically processed in this region corresponds to semantics. This finding is also consistent with the opinion that while the fundamental neural circuit for reading is universal, linguistic features in each language introduce different demands for specific cognitive components (Perfetti et al., 2013; Rueckl et al., 2015).

In addition to the left frontal cortices, a left parietal area also displayed neural integration in the “congruent < incongruent” direction and did not show additive enhancement, implying that higher activation in the incongruent condition is more likely to be a downstream effect. Unlike the left MFG, however, no activation was observed in either unimodal condition, and there was no integration-reading correlation. Therefore, although previous studies regarded the left parietal cortex as a part of the semantic network (Binder & Desai, 2011; Wang, Zhao, Zevin, & Yang, 2016), there was no evidence indicating the neural integration in this area reflects semantic processing. In another line of research, compared with English, the left parietal region was more recruited in Chinese reading for visual orthographic analysis given the visual complexity of characters (Perfetti et al., 2013). In this case, the neural integration effect might reflect a conflict of orthographic information. However, here we found no activation in the visual condition, going against this explanation. Third, although the left partial cortex has not been associated with print-sound integration previously in one’s native language, a recent training study revealed that this region was involved in acquiring novel grapho-phonological mapping (Xu, Kolozsvari, Oostenveld, & Hamalainen, 2020). As its function in audiovisual processing remains unclear, future studies are needed.

### Left STG: a potential link between pinyin-sound integration and oral reading fluency

In the pinyin experiment, no significant neural integration in either direction was observed. This is unexpected since, based on the previous studies in transparent scripts (Blau et al., 2010; van Atteveldt et al., 2004), stronger activation in the congruent condition than the incongruent condition should be found in the STCs. There are several possible reasons. First, the stimuli we used were at the syllable level. Differences between the letter-phoneme integration and string-syllable integration have been reported in previous studies (Kronschnabel et al., 2014). However, the authors also found that neural response to word-like stimuli is more sensitive to reading abilities. Thus, the alternative possibility is that the direction and strength of the neural integration are associated with reading abilities and masked by group averaging. In line with the latter, we did observe a positive correlation between the magnitude of the neural integration in the left STG and oral reading fluency. Specifically, children with higher oral reading scores are more likely to show stronger neural integration in the “congruent > incongruent” direction. In contrast, those with lower scores are more likely to show weaker neural integration or in the “congruent < incongruent” direction. This finding is consistent with studies that reported a reduced neural integration in individuals with RD in shallow languages (Blau et al., 2010; Blau et al., 2009), studies exploring brain-behavior correlation in both transparent (Wang et al., 2020) and opaque languages (McNorgan et al., 2014), as well as studies using a training paradigm (Karipidis et al., 2016; Karipidis et al., 2018).

The left STG is the central area representing phonological information (Boets et al., 2013; Glezer et al., 2016) and is involved in visual phonological processing in Chinese children and adults (Cao et al., 2010). Its activation and connectivity change after learning visual-sound mappings (Dehaene et al., 2010; Li, Xu, Luo, Zeng, & Han, 2020; Thiebaut de Schotten, Cohen, Amemiya, Braga, & Dehaene, 2014), which can further scaffold later reading development (Wang, Joanisse, & Booth, 2020). Noteworthily, the positive correlation between the neural integration in the left STG for pinyin-sound pairs and oral reading fluency was driven by a positive correlation in the congruent condition. This is different from that in the left MFG in processing character-sound pairs, indicating different mechanisms. It should also be noted that the current findings were revealed in typical readers. Whether children with RD have deficits or show other processing patterns requires further investigation.

The correlation between the neural integration in the left STG in processing pinyin-sound pairs with oral reading fluency also implies that learning pinyin may help children develop the underlying cognitive-linguistic components and corresponding neural circuits for fluent oral reading (though causal claims cannot be made from this study). Compared with cognitive-linguistic skills such as PA and RAN, less attention has been given to pinyin processing in understanding Chinese reading development, partially because of its auxiliary role as a scaffold in reading acquisition but not reading itself. Although the number of studies is limited, the pinyin-reading relationship has been demonstrated (Lin et al., 2010; Pan et al., 2011; Siok & Fletcher, 2001; Zhang, Georgiou, Inoue, Zhong, & Shu, 2020). Pan et al. (2011) found that invented pinyin spelling at age 6 independently predicted reading performance at ages 8 and 10 (Pan et al., 2011). Zhang and colleagues’ recent finding suggests a bidirectional relationship (Zhang et al., 2020).

Several explanations were proposed to explain the role of pinyin in reading development. One is that pinyin can help to establish better PA, the cognitive processing shared by pinyin and character reading (Ding, Liu, McBride, & Zhang, 2015; Li et al., 2020; McBride, Wang, & Cheang, 2018; Shu et al., 2008; Yin et al., 2011). Above and beyond PA, there is also a unique contribution from pinyin processing skills to character reading (Ju, Zhou, & delMas, 2021). Therefore, it is reasonable to hypothesize that pinyin may also help establish the correspondence between print and sound. In support of this notion, a study revealed that training on pinyin uniquely improved character-to-syllable mapping in Chinese as a second language learner (Guan et al., 2011). In the present study, a spatially overlapping region in the left STC also showed additive enhancement and neural integration in processing character-sound pairs. Furthermore, we found that the neural integrations in processing two stimuli categories were correlated. Given that the left posterior STC is involved in visual rhyming judgment of Chinese characters and proposed to play a role in mapping between orthography and phonology (Cao et al., 2010), the current findings further suggest that learning pinyin may shape the neural circuit for print-sound integration that will later be recruited in learning characters.

### Caveats and future directions

This study investigated the neurofunctional basis of integrating audiovisual information of characters and pinyin, respectively. While it represents one of the first efforts in this field, caution should be considered when interpreting the results. (a) This study focused on print-sound integration at the syllable level, whereas previous studies mainly explored letter-speech sound associations (but see Kronschnabel et al., 2014; Wang et al., 2020). The primary reason is that there are no GPC rules in the Chinese writing system, and print-sound mapping occurs at the syllable level. Of importance, single characters are taught at the earliest stages and play fundamental roles in reading development (Shu et al., 2003; Xing, Shu, & Li, 2004), similar to the role of GPC rules in alphabetic languages. (b) Given that our participants were in grades 3-5, pinyin may play a different role than it does in the earlier stages of reading development. For example, Siok and Fletcher (2001) unveiled significant correlations between pinyin knowledge and reading in grades 2-5, but not in grade 1. Thus, it could be the age range of the children in this study that enabled us to observe the relationship between print-sound processing in pinyin and oral reading fluency. To further answer this question, it is necessary to conduct studies following preliterate children on the pinyin and reading development until they achieve fluent reading. (c) In the present study, the pinyin stimuli were always presented before the characters to prevent priming. To investigate script-specific activation and brain-reading correlation while considering the order effect, future research could increase the sample size and assign the experiment orders counterbalanced. (d) In the voxel-wise whole-brain analysis, we used a liberal threshold (*p* < 0.005) at the voxel level and applied the FWE correction to control Type I error (α < 0.05) at the cluster level. While the voxel lenient threshold has been common in pediatric research (Chyl et al., 2018; Plewko et al., 2018), it may inflate false positives (Eklund, Nichols, & Knutsson, 2016). Therefore, studies with larger sample sizes are needed to replicate the current findings in the future. (e) The previous studies have revealed reading predictors (e.g., phonological processing, language skills) at the preliterate stage and the corresponding neural substrates (Marks et al., 2019; Saygin et al., 2013), which could be the element of print-sound integration. It is necessary to depict the whole picture by conducting longitudinal research that includes all these measures. (f) Finally, here we uncovered the brain bases of print-sound integrations in two contrasting scripts and their relationship with reading performance in typically developing children. One next question that warrants investigation in the future is whether neural processes are altered in children with RD.

## Conclusion

In the current study, we simultaneously investigated the neural mechanisms underlying the audiovisual integration of character-sounds and pinyin-sounds in a non-alphabetic language. The results revealed additive enhancements in the bilateral STCs as a universal brain feature accompanying audiovisual information processing across scripts. Moreover, the precise manifestation of neural integration is influenced by linguistic features. In particular, the left MFG is involved in the audiovisual integration of characters and correlated with reading comprehension, suggesting that semantic representations could be more efficiently accessed and processed in readers with higher comprehension proficiency. On the other hand, the brain-behavior correlation and brain-brain correlation in the left STC imply that learning pinyin may help children develop better reading skills by promoting the development of the shared cognitive-linguistic component and shaping the underlying brain circuits.

## Acknowledgments

The authors would like to thank all the children and their families for participating in this study and all the testers for helping with data collection. This work was supported by the Key Program of the National Social Science Foundation of China (14ZDB157), National Key Basic Research Program of China (2014CB846103), National Natural Science Foundation of China grants (31271082, 31671126, 31611130107, 61374165, 81801782), Beijing Municipal Science & Technology Commission (Z151100003915122), Fundamental Research Fund for the Central Universities, China Postdoctoral Science Foundation (2019T120062, 2018M641235), and Social Science Fund of Beijing (17YYA004).

## Author Contributions

Conceptualization: ZX, HS, XL; Investigation: ZX, TY, XC, HL, XZ; Formal Analysis: ZX; Data Curation: ZX; Writing – Original Draft Preparation: ZX; Writing – Review & Editing: ZX, TY, XC, FH, HL, XZ, HS, XL; Funding Acquisition: ZX, HS, XL; Supervision: ZX, HS, XL

## Data Availability Statement

The data that support the findings of this study are available from the corresponding authors upon reasonable request. The data are not publicly available due to privacy or ethical restrictions.

## Abbreviations

ANOVA: analysis of variance
Aud: auditory
avC: audiovisual congruent
avI: audiovisual incongruent
fMRI: functional magnetic resonance imaging
FWE: Family-Wise Error
GPC: grapheme-phoneme correspondence
IQ: intelligence quotient
MEG: magnetoencephalography
MFG: middle frontal gyrus
MNI: Montreal Neurological Institute
PA: phonological awareness
RAN: rapid naming
MA: morphological awareness
RD: reading disorder
ROI: region-of-interest
RT: reaction time
SD: standard deviation
STC: superior temporal cortex
STG: superior temporal gyrus
Vis: visual
TR: repetition time
WISC-CR: Chinese Wechsler Intelligence Scale for Children

