## Supplemental Materials for "Neurofunctional basis underlying audiovisual integration of print and speech sound in Chinese children"

**Table S1** In-scanner performance

| | Overall | Run 1 | Run 2 | *t*-test | Vis | Aud | avC | avI | ANOVA | Post-hoc Test |
|---|---|---|---|---|---|---|---|---|---|---|
| *Character* | | | | | | | | | | |
| ACC | 0.965 (0.048) | 0.957 (0.068) | 0.970 (0.042) | $t = -1.604$ $p = 0.119$ | 0.968 (0.063) | 0.934 (0.110) | 0.972 (0.061) | 0.983 (0.044) | $F = 1.002$ $p = 0.395$ | --- [a] |
| RT (ms) | 512 (81) | 516 (87) | 507 (86) | $t = 0.962$ $p = 0.344$ | 575 (79) | 531 (119) | 461 (80) | 482 (89) | $F = 35.819$ $p < 0.001$ | Vis > Aud > avC = avI |
| *Pinyin* | | | | | | | | | | |
| ACC | 0.964 (0.037) | 0.965 (0.046) | 0.963 (0.042) | $t = 0.223$ $p = 0.825$ | 0.913 (0.105) | 0.966 (0.074) | 0.985 (0.050) | 0.988 (0.038) | $F = 10.067$ $p < 0.001$ | Vis < Aud = avC = avI |
| RT (ms) | 530 (80) | 527 (75) | 531 (92) | $t = -0.599$ $p = 0.553$ | 601 (90) | 566 (125) | 471 (67) | 496 (76) | $F = 54.077$ $p < 0.001$ | Vis > Aud > avC = avI |

*Note*. Standard deviations are presented in the parentheses. [a] No post-hoc test was conducted given that the main effect of conditions was not significant. Abbreviations: ACC = accuracy, RT = reaction time, Vis = unimodal visual, Aud = unimodal auditory, avC = audiovisual congruent, avI = audiovisual incongruent.

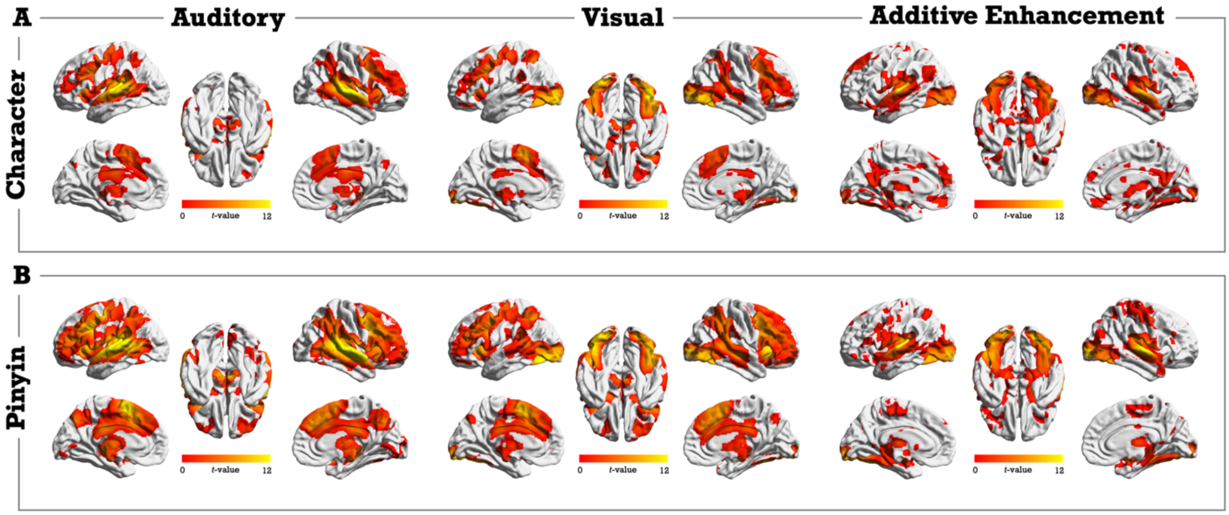


**Figure S1** Brain maps of activation in the unimodal auditory and visual conditions, as well as the additive enhancement (audiovisual congruent > the average activation of auditory and visual conditions) in the character (A) and pinyin (B) experiments. The uncorrected threshold of *p*-voxel < 0.05 was used. The three brain maps for one script were used to identify regions showing audiovisual additive enhancement with a conjoint *p*-voxel < 0.05^3 = 0.000125.
